# Encoding sensing functions into material interface for a rationally engineered integrated electrochemical liquid biopsy

**DOI:** 10.1101/2022.01.17.476350

**Authors:** Yuan Zhang, Hao Zhu, Zi Ying, Xinghua Gao, Wei Chen, Yueping Zhan, Lingyan Feng, Chung-Chiun Liu, Yifan Dai

## Abstract

Limited healthcare capacity highlights the needs of integrated and simple sensing systems for personalized health monitoring. However, only a limited set of sensors can be employed for point-of-care applications, emphasizing the lack of a generalizable engineering strategy for sensor construction. Here, we report a *de novo* rational engineering strategy for the construction of an integrated electrochemical liquid biopsy (ELB) platform capable of direct profiling cancer exosomes from blood. Using a bottom-up approach for sensor design, a series of critical sensing functions is considered and encoded into the material interface by programming the electrode material with different chemical and structure features. We present that the rationally engineered electrochemical liquid biopsy platform is able to achieve one-step sensor fabrication, target isolation, non-fouling and high-sensitivity sensing, direct signal transduction and multiplexed detection. Integrating the multiplexed sensing with principal component analysis, we demonstrate the capability of the programmed sensing system on differentiating cancerous groups from healthy controls by analyzing clinical samples from lung cancer patients.

## Introduction

The ongoing severe acute respiratory syndrome coronavirus 2 (SARS-CoV-2) pandemic poses an imminent healthcare challenge to the world. The high infectivity, long incubation time and asymptomatic nature of SARS-CoV-2 result widespread transmission across the world with over 310 million cases^1^. As a result, current healthcare capacities are overwhelmed across the world, significantly impacting the timely diagnostics and treatments of patients with other types of severe diseases^2^, especially metastatic cancers^3^. Such unexpected scenarios highlight the long-term needs of simple, reliable, and accurate diagnostic systems for metastatic cancer management.

Liquid biopsy, which detects tumor biomarkers directly from blood sample, is the current gold-standard for non-invasive metastatic cancer managements^4^. However, the analytical methods applied in liquid biopsy include enzyme-linked immunosorbent assays (ELISA) for protein analysis and next-generation sequencing for nucleic acid analysis^5^, which require laboratory settings, technical expertise and long turnaround time to achieve reliable diagnostic, limiting its applicability in resource-limited areas, especially in current unusual time.

To tackle the challenges for the developments of a simple, quantitative, and accurate assay toward detecting metastatic cancers, we herein present a *de novo* engineering strategy for the development of electrochemical liquid biopsy (ELB) platform, enabling direct sorting and multiplexed analysis of different biomarkers on tumor-derived extracellular vesicles. By encoding critical sensing functions into the material interfaces, the developed ELB is capable of direct quantitative detection of cancers from a single drop of blood within 30 min. ELB was employed for the analysis of blood from patients with lung cancer, showing excellent correlation with clinically established cancerous group. Owing to the modularity of our demonstrated engineering approach, the ELB can be programmed to a generalizable platform technology for the detection of other different biomarkers.

## Results

### Rational engineered hybrid nanostructured electrode as a sensing interface

For the development of multiplexed, point-of-care ELB, we rationally evaluated the technical criteria that are imperative to enable a fully integrated ELB system (**Fig. 1**). Specifically, for the purpose of direct analysis of cancer related extracellular vesicles, the following sensing features should be encoded into the sensing interface: 1) efficient electron transfer ensuring high signal-to-noise ratio to achieve an ideal detection sensitivity (i.e. detection limit) and sensing resolution (i.e. signal differentiation), 2) specific target sorting from complex biological samples, 3) non-fouling surface ensuring detection sensitivity in complex biological matrix, 4) direct target sensing with limited operational procedures, 5) multiplexed analysis guaranteeing the high-accuracy of the platform. To achieve these desired functions, we *de novo* designed and constructed a miniaturized and mass-producible nano-structured sensor system through programming the material properties of the hybrid nanocomposites.

**Figure. 1.**
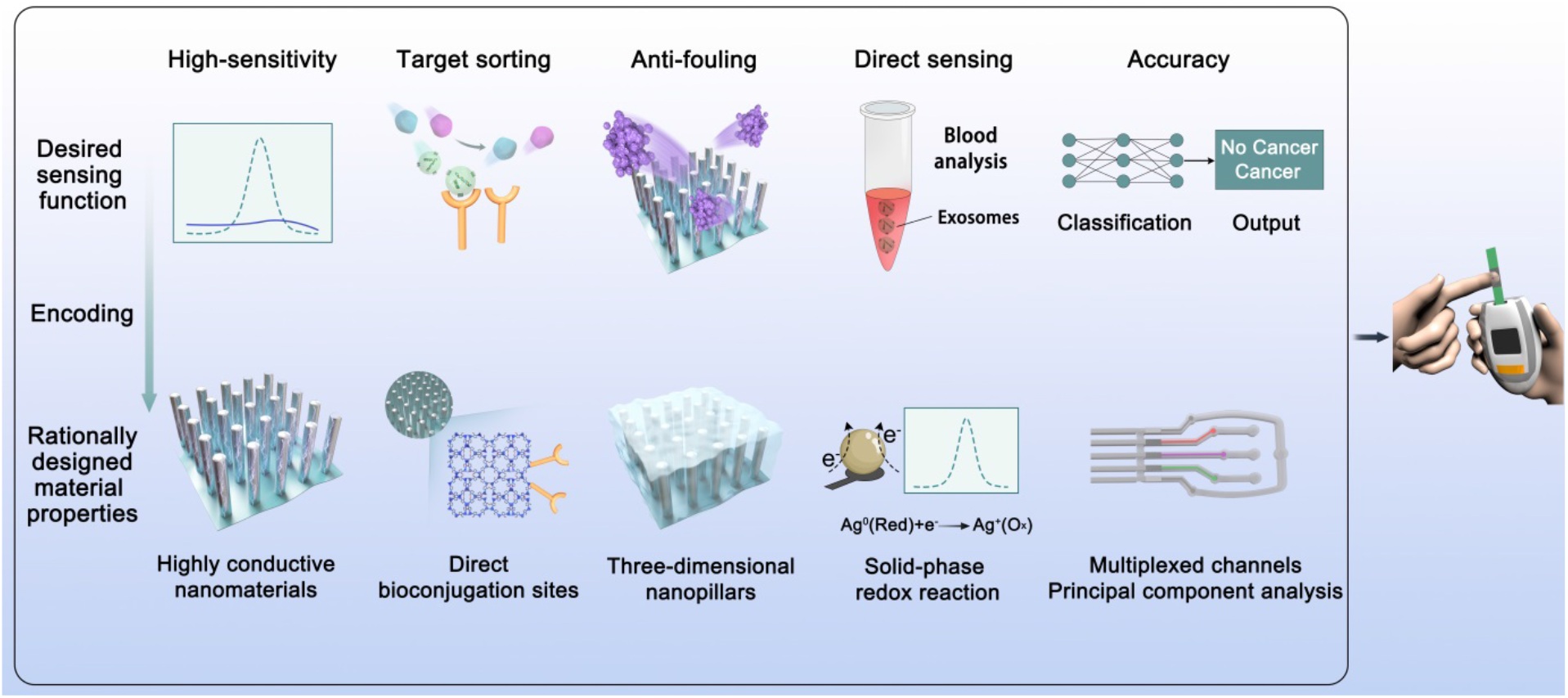
Schematic illustration of the rational engineering strategy toward the development of electrochemical liquid biopsy. Desired sensing functions serve as the criteria for the design of sensing interface. Through bottom-up programming and constructing the material properties of the sensing interface, functional sensing features are encoded. This rational engineering process delivers a fully integrated electrochemical liquid biopsy platform technology.

To construct a sensing interface enabling a high signal-to-noise ratio, we employed a nanostructured pillar array providing efficient mass transport and electron transfer channels for signal amplification^6^. Specifically, ordered ZnO nanopillar was grown onto ultrathin 1T MoS_2_ nanosheets forming a hybrid nanostructure. In order to achieve exosomes separation from blood, we applied a ZIF-90 thin film containing aldehyde group (-CHO) for bioconjugation to immobilize recognition antibodies of exosome proteins through direct interacting with -NH_2_ groups on the antibodies. In the ZIF-90 thin film, abundant–CHO groups are periodically arranged in the rigid metal-organic framework^7^, avoiding the potential degradation of antibody by using conventional flexible glutaraldehyde molecule^8^. Target bioconjugation can induce the desired conformation of antibody^9,10^, and would be in favor of the maximized capture efficiency of exosomes through affinity binding. Furthermore, such three-dimensional (3D) porous matrix, by simply interlacing with bovine serum albumin (BSA), can achieve anti-fouling property^11^. To achieve direct quantification, we took advantages of the customized electrodes, in which solid-phase electroactive species (silver/carbon composite) were paved underneath the nano-structured electrode for redox reaction^12^. Therefore, with the presence of the highly negatively charged exosomes^13^ on the surface, the electron transfer process would be significantly hindered, realizing quantitative difference in electrochemical current. Comparing with conventional electrochemical quantification methods through investigating the change of surface impedance^14^, by encoding the redox reaction into the materials, we eliminated the use of typical electrochemical redox molecules^14^ (e.g. ferricyanide or ruthenium nitride) which can potentially cause irritation of eyes and skins, providing a more user-friendly and safe sensing system. Finally, to fulfill the need of high-accuracy exosomes analysis, we designed a three-channel microfluidic sensor chip, enabling simple preparation process of ELB and multiplexed simultaneous sensing. To test the capability of this platform, we selected CD9, CD63 and CD81^15,16^ as the detecting targets, which residue on the surface of the cancer related extracellular vesicles and have been clinically validated for their efficacy. To prevent false-positive/negative results, we employed principal component analysis to process the electrochemical data, maximizing sample variations.

### Design of microsensor chip enables direct signal transduction

The microsensor chip is composed of screen-printed electrodes sealed with polydimethylsiloxane (PDMS) layer. Five electrodes, including counter electrode (CE), reference electrode (RE) and three different working electrodes (WE), are printed on a polyethylene terephthalate (PET) substrate (See methods for details, **Supplementary Fig. 1**). To enable direct electrochemical sensing without external introduction of redox active chemicals, we first evaluated whether electrochemical redox reaction can be encoded into the solid-phase materials. We evaluated whether the embedded silver/carbon electrical conduction band underneath the WE is capable of generating electrochemical signal, owing to the known Ag^0^/Ag^+^ redox reaction^17,18^. By applying cyclic voltammetry (CV) in phosphate buffer, a pair of redox peaks corresponding to consecutive one-electron-transfer reactions of Ag^0^(Red)/Ag^+^(Ox) (E_Ox_=0.08 V vs. Ag/AgCl) and Ag^+^(Ox)/Ag^0^(Red) (E_Red_=-0.13 V vs. Ag/AgCl) was observed (**Supplementary Fig. 2**), confirming our capability to encode redox reaction into solid-phase materials.

### Characterizations of rationally engineered nanostructured sensing material

Based on the considerations set for an ideal sensing interface toward electrochemical analysis, we designed and engineered the metal organic framework MoS_2_/ZnO/ZIF-90 based nanostructure as the materials of working electrode through hierarchical growth (see Methods) to achieve robust electron transfer and low-matrix effect^11^. Material and surface characterizations were performed to evaluate the chemical identity and structure of the designed material.

First, SEM images of MoS_2_/ZnO sample show that the ordered nanopillars array is uniformly grown onto ultrathin MoS_2_ nanosheets (**Fig. 2a**). The high-magnification of SEM image reveal that the diameter of vertical ZnO nanopillars is 20-40 nm and a glazed surface is observed over nanopillars (**Fig. 2b**). After exposure to imidazole-2-carboxaldehyde (2-ICA) vapor, the surface of ZnO nanopillars became roughened with bumpy and hollow structures (**Fig. 2c**). Elemental mapping analysis in nanopillars region demonstrates the homogeneous distribution of Zn, O, C and N species **(Fig. 2d**). It should be noted that the contents of C and N are slightly lower than that of another two species, indicating that only the surface portion of ZnO nanopillars was turned into ZIF-90. The low-magnification TEM image shows that the length of nanopillars is about 1.5 μm (**Fig. 2e**), and further HRTEM characterization demonstrates that the nanopillars are core-shell structure with 8-12 nm of ZIF-90 layer formed on the surface (**Fig. 2f**). During the growth of ZIF-90 thin layer, we found the reaction time is critical for the formation and structure. With optimized reaction time, the integrated MoS_2_/ZnO/ZIF-90 nanoarchitecture is successfully prepared (**Supplementary Fig. 3**).

**Figure. 2.**
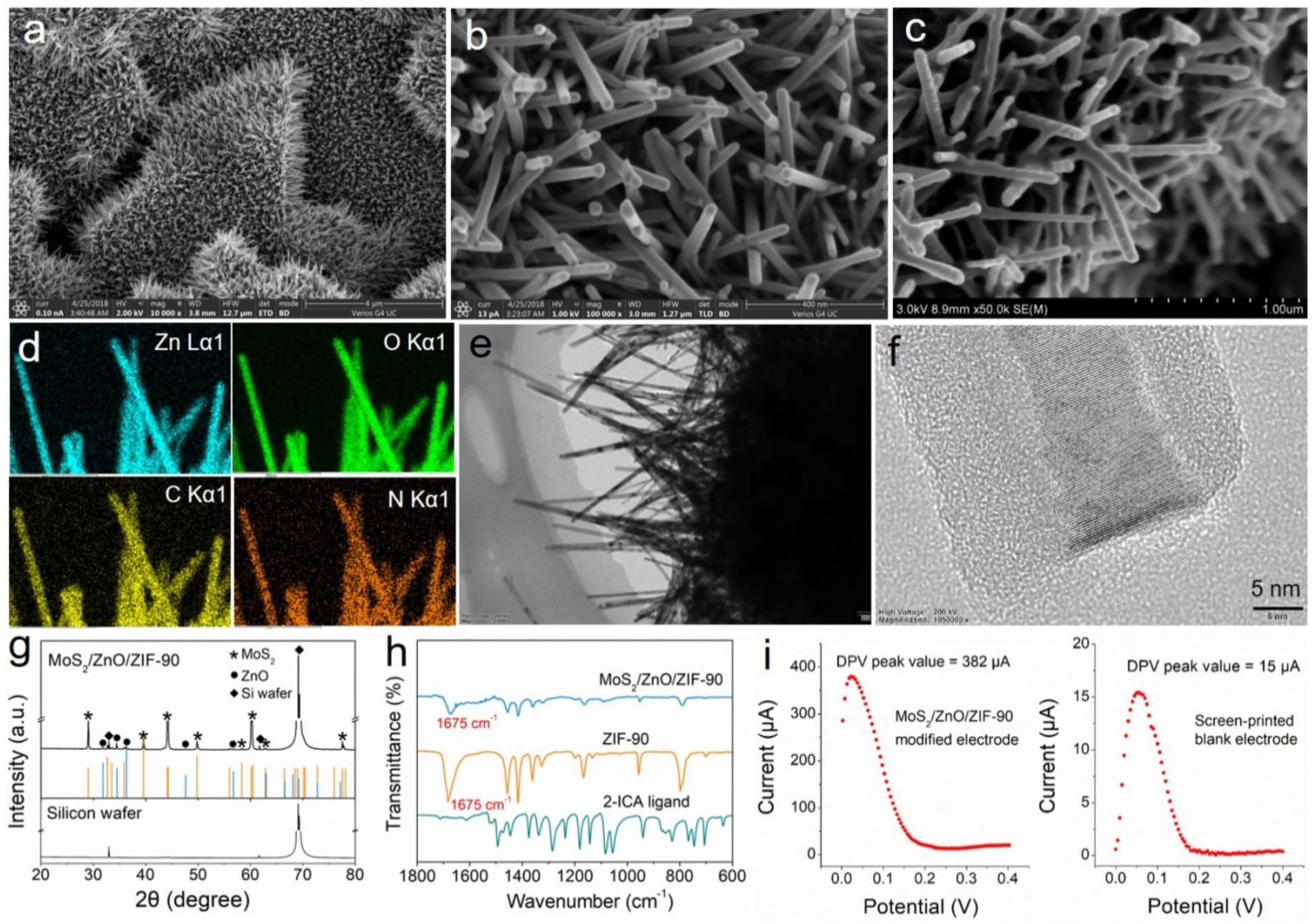
Characterizations of MoS_2_/ZnO and MoS_2_/ZnO/ZIF-90 nanostructures based working electrode and the configuration of fabricated microsensor. **a, b**, Low-magnification and high-magnification SEM images of MoS_2_/ZnO. **c**, High-magnification SEM image of MoS_2_/ZnO/ZIF-90. **d**, Elemental mapping analysis of MoS_2_/ZnO/ZIF-90. **e**, TEM image of MoS_2_/ZnO/ZIF-90. **f**, HR-TEM images of MoS_2_/ZnO/ZIF-90. **g**, XRD pattern of MoS_2_/ZnO/ZIF-90. **h**, FT-IR spectra of MoS_2_/ZnO/ZIF-90, ZIF-90 and imidazole-2-carboxaldehyde. **i**, Programmed materials mediated amplification of signal amplification as demonstrated through measurements by differential pulse voltammetry on MoS_2_/ZnO/ZIF-90 modified or unmodified screen-printed electrode in PBS solution (pH=7.2).

Furthermore, to evaluate the morphological feature of the complex nanostructure, we applied electron microscope tomography. The micrographs of MoS_2_/ZnO/ZF-90 sample were recorded from various orientations by tilting the specimen, which were then computationally merged into a 3D structure (**Supplementary Fig. 4**). According to these images, ultrathin MoS_2_ material displayed as blue color is located at the bottom of the 3D nanoarchitecture, and the outermost green is originated from ZIF-90 thin film. Additionally, the red ZnO nanopillars are appeared to be orange due to its overlapping with the outer green thin film. These electron microscopy 2D and 3D images provide the detailed structure information on the integrated nanoarchitecture.

Moreover, X-ray diffraction (XRD) characterization shows the pattern consistency between the fabricated MoS_2_ with standard 2H-MoS_2_ structure (JCPDS No. 03-065-0160), and the peaks labeled with black dots are corresponding to the strongest peaks of ZnO (JCPDS No. 01-075-0576) (**Fig. 2g**). No peak from ZIF-90 was observed because only a thin layer of ZnO is consumed and transformed into ZIF-90. Fourier-transform infrared spectroscopy (FT-IR) was then performed (**Fig. 2h**) and all the peaks in the spectrum of MoS_2_/ZnO/ZIF-90 sample are in excellent agreement with that of pure ZIF-90 and no characteristic peak derived from 2-ICA ligand is present, indicating the successful transformation of 2-ICA to ZIF-90 instead of simple surface absorption. The bands in the spectral region of 600-1500 cm^-1^ are associated with the stretching and bending modes of the imidazole ring^19,20^. Most importantly, the characteristic peak at about 1675 cm^−1^ reveals the presence of –CHO in materials^21,22^, confirming our capability on direct immobilization of recognition elements. These characterizations confirmed the chemical identity and structure of the programmed sensing materials.

To test whether such nanostructured electrode enhances electron transfer, we applied differential pulse voltammetry (DPV) to compare the electrochemical signal before and after the addition of MoS_2_/ZnO/ZIF-90 layer. The peak current of modified electrode leads to a 2446.7% increase comparing with unmodified electrode (**Fig. 2i**), confirming capability of signal amplification by the developed nanostructured electrode^23,24^.

### Microfluidic channels enable simple sensor fabrication and multiplexed sample analysis

To achieve multiplexed analysis, we set out to integrate multi-channel microfluidics as the sample inlets for the electrochemical microsensor **(Fig. 3a**). The microchannels on the chip can be employed for the immobilization of bio-recognition antibodies for sensor fabrication as well as simultaneous multiplexed detection of exosome markers.

**Figure 3.**
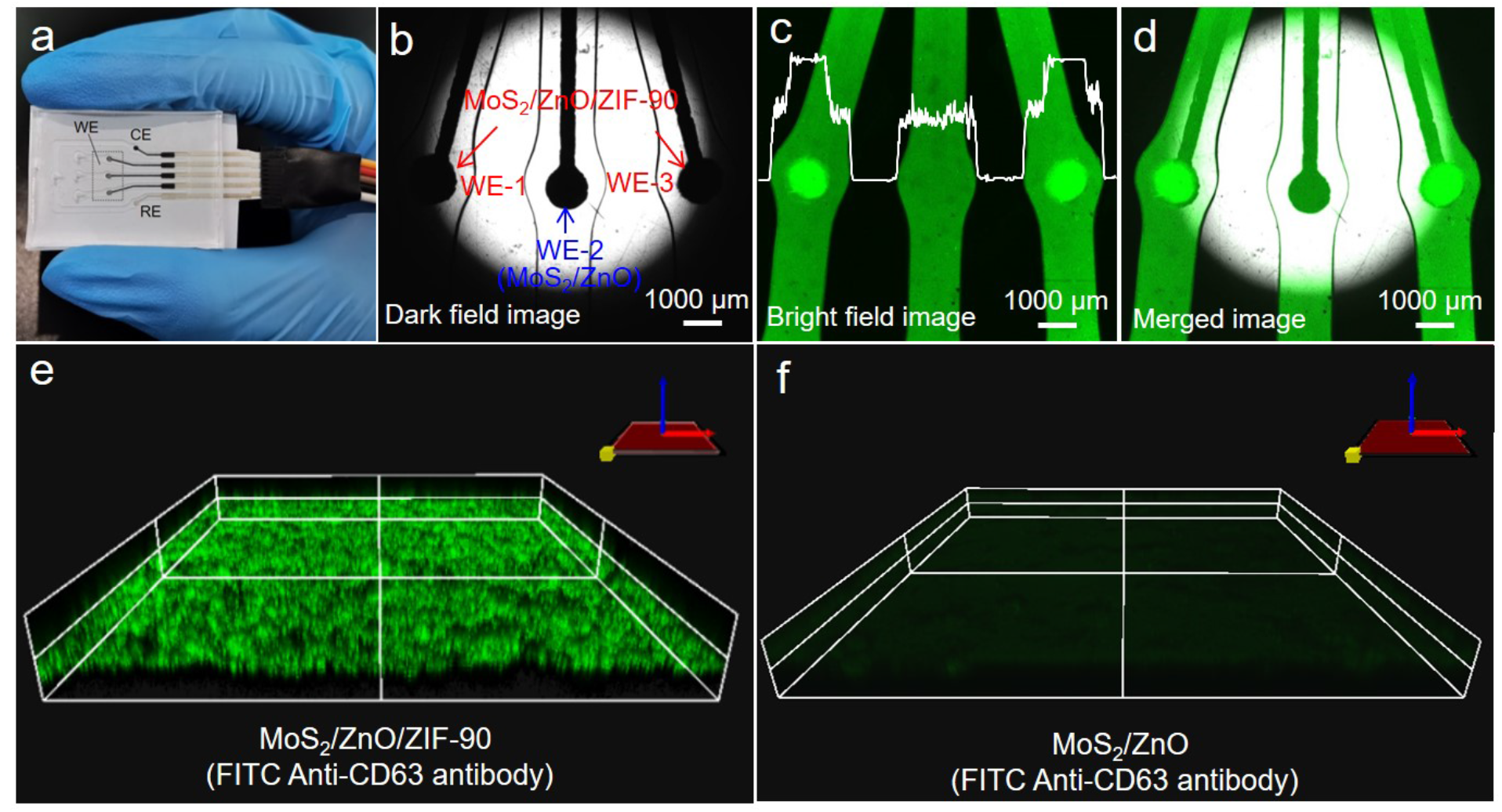
Direct immobilization of recognition elements onto the nanopillar. **a**, An integrated three-channel sensor chip (WE=working electrode, CE=counter electrode and RE=reference electrode). **b, c, d**, Dark field, bright field and merged fluorescence images of different WEs. Two of WEs (WE-1 and WE-3) are deposited with MoS_2_/ZnO/ZIF-90 nanohybrids, and WE-2 is loaded with MoS_2_/ZnO nanomaterial. All three WEs are incubated with FITC Anti-CD63 antibody. **e, f**, 3D fluorescence images of MoS_2_/ZnO/ZIF-90 and MoS_2_/ZnO after immobilized with FITC Anti-CD63 antibody.

It is critical to ensure the sealing ability and orthogonality of microchannels for processing liquid samples. To test this notion, solutions containing different dyes were added into the microchannels and no leakage was observed (**Supplementary Fig. 5**). We further introduced different dyes separately into three microchannels from their corresponding inlets, demonstrating orthogonality on liquid transportation. Therefore, these channels can be directly applied to immobilize Anti-CD9, Anti-CD63 and Anti-CD81 antibodies onto WE-1, WE-2 and WE-3 respectively, presenting a simple procedure for sensor fabrication (See methods for details and **Supplementary Fig. 1**).

For the purpose to achieve direct multiplexed analysis of sample, a fourth inlet was designed to accommodate testing sample, allowing flow diversion into all the WEs. To ensure sensing accuracy, we tested the consistency of output signals through different microchannels. The phosphate buffered saline (PBS) solution (pH= 7.2, 100 μL) was filled into the chip channels, and the obtained CV curves overlap with each other (**Supplementary Fig. 6**), indicating that the engineering and fabrication techniques are reliable to produce reproducible sensing interface.

### Characterizations of the ZIF-90 for immobilization of recognition elements

For the purpose of target sorting and detection of exosomes, the prepared MoS_2_/ZnO/ZIF-90 material was tested for antibody immobilization and the following immunoreaction. A fluorescein isothiocyanate (FITC) labeled antibody was employed and the confocal fluorescence microscope was used to observe the results of immobilization. Here, FITC Anti-CD63 is served as an illustration. We applied WE-1 and WE-3 pre-deposited with MoS_2_/ZnO/ZIF-90 and WE-2 (located in the middle of the sensor chip) pre-deposited with only MoS_2_/ZnO material for the immobilization of FITC Anti-CD63. Comparing the dark field, bright field and merged fluorescence images of different WEs (**Fig. 3b-d**), two clearly defined spots were noticed in the region where the electrodes are covered by MoS_2_/ZnO/ZIF-90 material, while no change in fluorescence intensity was observed along the middle channel. The 3D fluorescence images also show that the uniform green fluorescence is appeared over MoS_2_/ZnO/ZIF-90 material, while there is almost no fluorescence observed by using MoS_2_/ZnO as electrode material (**Fig. 3e-f**). Moreover, a vertically array structure is dimly visible in the whole observation area, further revealing that FITC Anti-CD63 is immobilized on the surface of nanopillars, potentially enhancing the target capture capability with such 3D structure^25^. These fluorescence characterizations confirm that MoS_2_/ZnO/ZIF-90 nanostructure could be used for the effective immobilization of antibody depending on the surface ZIF-90 thin film. To verify the antibody immobilization is solely attributed to ZIF-90 instead of a cooperative effect of the nanostructures, we employed ZIF-90 nanoparticles for antibody immobilization (**Supplementary Fig. 7**). Similarly, strong fluorescence intensity is also detected over ZIF-90 nanoparticles. Although pure ZIF-90 nanoparticles possess the ability to effectively immobilize recognition antibody, only a minute current response was measured (**Supplementary Fig. 8**). This control experiment illustrates the importance of the synergistic effect of the sensing materials on sensing functions.

### EIS confirms the principles of the nanostructured electrodes toward high-sensitivity analysis and immunoreactions

The electrochemical impedance spectroscopies (EIS) of different materials as well as the developed microsensor for immunoassay were measured (**Fig. 4a**) and displayed with an impedance order of silver/carbon screen-printed electrode, ZIF-90, MoS_2_, MoS_2_/ZnO/ZIF-90, MoS_2_/ZnO/ZIF-90-Anti CD63 (MoS_2_/ZnO/ZIF-90-Ab), MoS_2_/ZnO/ZIF-90-Anti CD63-blocking buffer (MoS_2_/ZnO/ZIF-90-Ab-BB), and MoS_2_/ZnO/ZIF-90-Anti CD63-blocking buffer-CD63 protein (MoS_2_/ZnO/ZIF-90-Ab-BB-Ag). Generally, EIS is an important electrochemical technique based on the interfacial reaction at the electrode surface, which could be used to evaluate the electron transfer resistance (R_et_) on electrode surface^26,27^. The EIS result is often represented by the Nyquist plot, including a semicircle at high-frequency and linear part at low-frequency region. Since the diameter of semicircle in Nyquist plot is related to R_et_^28,29^, the EIS curve of MoS_2_ shows the disappearance of semicircle at high-frequency region compared with silver/carbon screen-printed electrode, demonstrating that the electron transfer rate is enhanced with the modification of MoS_2_. However, a clear semicircle was obtained over ZIF-90 modified electrode, which reveals the large R_et_ of ZIF-90 particles. After modification of MoS_2_/ZnO/ZIF-90 material, only the linear part is appeared and the linear slop is higher than that of pure MoS_2_, which indicates that the hybrid nanostructure is more favorable for electron transfer than 2D MoS_2_ nanosheets. The Nyquist graphs and fitted values show that the most significant change occurs in R_et_, and the R_et_ values of these systems are 61.78 Ω (screen-printed electrode), 156.70 Ω (ZIF-90 modified electrode), 34.43 Ω (MoS_2_ modified electrode) and 15.73 Ω (MoS_2_/ZnO/ZIF-90), respectively. These results also suggest that the developed nanostructure with an excellent electron transfer capability is beneficial to signal amplification for our sensor system.

**Figure 4.**
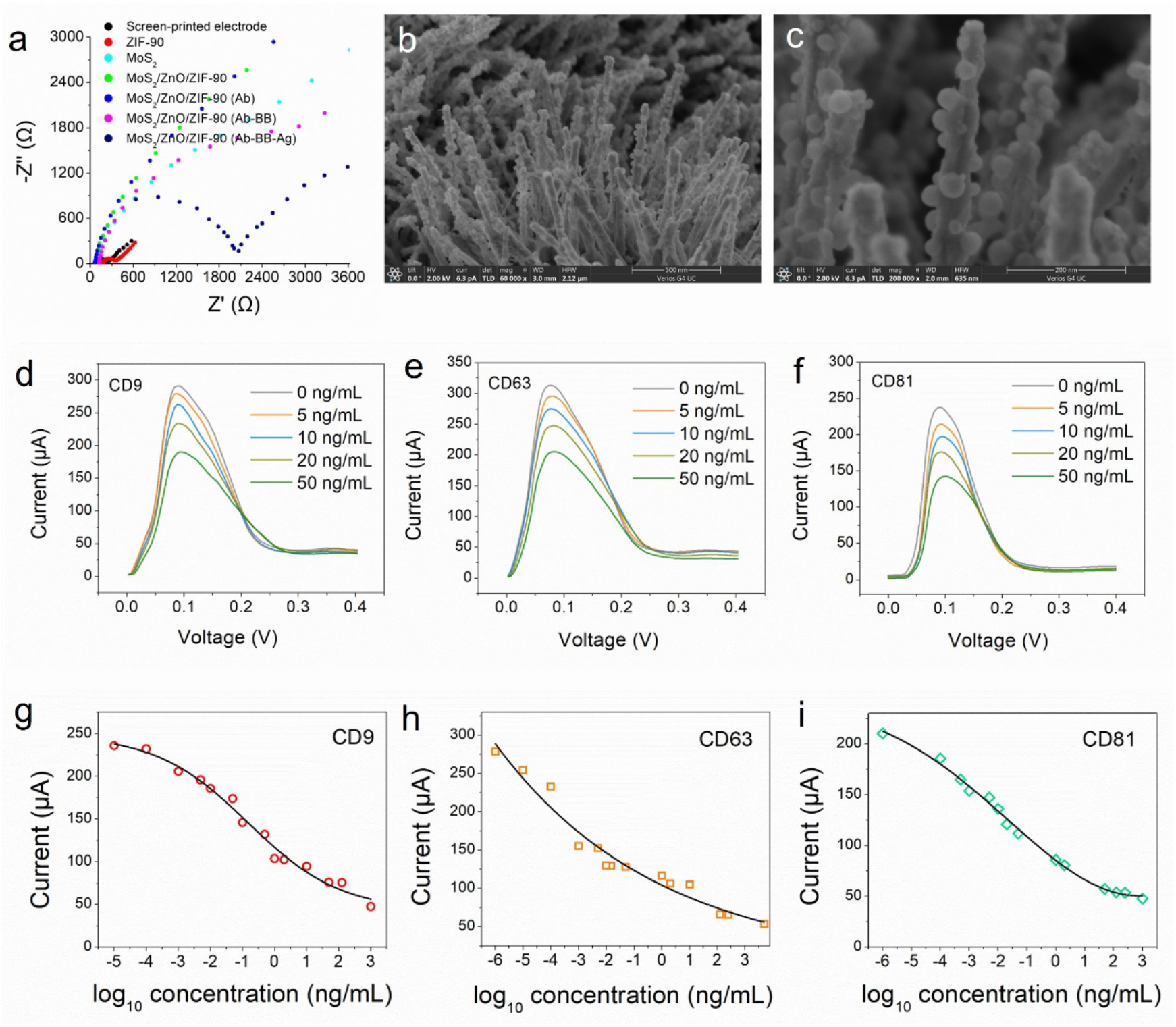
Characterizations of the programmed materials on target sorting and quantitative sensing. **a**, EIS measurement (Nyquist plots) of different materials (ZIF-90, MoS_2_ and MoS_2_/ZnO/ZIF-90) as well as the developed microsensor during the preparation process. **b**, SEM image showing the uniformly captured exosomes on the surface nanopillars. **c**, High-magnification SEM image revealing the size and morphology of exosomes. **d, e, f**, DPV measurements of the developed microsensor in the presence of CD9 (WE-1), CD63 (WE-2) and CD81 (WE-3) at various concentrations. **g, h, i**, Dose-dependent responses of exosomes from culture supernatant based on the detection of CD9, CD63 and CD81.

Followed by incubation of antibody with MoS_2_/ZnO/ZIF-90, the Nyquist plot appears limited variation, due to the low incubation concentration and neutral charge property of the antibodies used. However, once we blocked the electrode surface with BSA, a depressed semicircle of Nyquist curve was observed. After the final immunoreaction, an obvious semicircle at the high frequency range emerged (**Fig. 4a**), implying the capability of the developed microsensor on electrochemical immunodetection of exosomes biomarker. The distinct semicircle from the EIS curve of MoS_2_/ZnO/ZIF-90-Ab-BB-Ag is ascribed to the formation of antibody-antigen complex which impedes the electron transfer on the surface of the electrode. From the above observations, we demonstrated that MoS_2_/ZnO/ZIF-90 nanohybrids not only possess the capability of enhancing electron transfer, but also feasible for immobilization of antibody as well as the following antigen–antibody interaction.

### Direct capture of extracellular vesicles from complex matrix

We then verified the exosomes sorting ability of the microsensor chip through antibody-antigen affinity reaction. By introducing complex biological matrix containing extracellular vesicles onto the sensor, in the presence of a recognition antibody (Anti-CD63), all the nanopillars are uniformly surrounded with exosome particles (**Fig. 4b)**, demonstrating effectiveness of our platform for exosomes capture. The high-magnification SEM image reveals that the diameter of captured exosomes is about 15 nm to 50 nm (**Fig. 4c**). Furthermore, the exosomes are enriched on the surface of the nanopillars, presenting a direct confirmation of that the significantly decreased electron transfer kinetics (**Fig. 4a**) were due to the presence of negatively charged vesicles.

### Verification of electrochemical sensing principle

For the multiplexed measurement of exosomes markers, we separately introduced different antibodies (Anti-CD9, Anti-CD63 and Anti-CD81) into the corresponding microchannels to construct the immunoelectrodes. Recombinant CD9, CD63 and CD81 proteins with different concentrations (prepared in PBS buffer) were added into three microchannels and incubated at 37 °C for 30 min. The DPV current responses of different exosomes markers decrease along with the concentrations (**Fig. 4d-4f**), ascribing to the formation of antibody-antigen complexes which block the electron transfer between the WE and CE. These results demonstrate that the engineered microsensors chip is capable of quantitative analysis exosomes markers. Additionally, the change of current signals due to CD9 and CD63 proteins are higher than that of CD81 protein, indicating that the current response caused by immunoreaction is related to the type of exosomes marker.

### Direct exosome quantification from cultures of human non-small cell lung cancer

We challenged our platform for the detection the exosomes from culture supernatant of human non-small cell lung cancer cell line A54. Bicinchoninic Acid (BCA) Protein Assay Kit was used for determining the concentration of exosomes extracted from cell supernatant (see Methods), and the measured concentration was 100 μg/mL. For quantification, different concentrations (5×10^4^ ∼ 1×10^−5^ ng/mL) of exosomes were obtained by dilution and added into the microsensors chip. The dose-dependent current signals were measured **(Fig. 4g-4i**), showing that the output signal was correlated with the concentration of exosomes. The developed microsensors chip can effectively detect exosomes in the concentration range from ng/mL to pg/mL. The limit of detection (LOD) was evaluated to be 10 fg/mL for CD9, 1 fg/mL for CD63 and 1 fg/mL for CD81, respectively. The LOD was superior to that of previous reports^30-33^. These results confirm the capability of the microchip to probe exosomes in complex samples.

### Analyzing human lung cancer blood samples

To demonstrate the practical application of our developed microsensor chip, human blood samples were analyzed. Twenty-four clinical blood samples were collected, twelve of them were from healthy individuals (H1-H12) and the other twelve were samples of cancer patients (P1–P12). All the samples were diluted to 15-fold in PBS solution and the current signal generated by blank PBS solution was also quantified as an experimental control. We found that the current response of clinical blood samples from healthy individuals was higher than the current from cancer patients (**Fig. 5a, b**), which demonstrates that exosomes from cancer patients express higher levels of CD9, CD63 and CD81 than healthy subjects. Specifically, the electrodes immobilized with Anti-CD63 and Anti-CD81 could effectively distinguish the cancer patients and healthy controls, while three of cancer samples and one in healthy group displayed the abnormal expression in the testing results of CD9. Since surface proteins on exosomes carry information about their tissues of origin, they may have different expression profiles on exosomes, thus inducing some variation in quantification of exosomes^15^.

**Figure 5.**
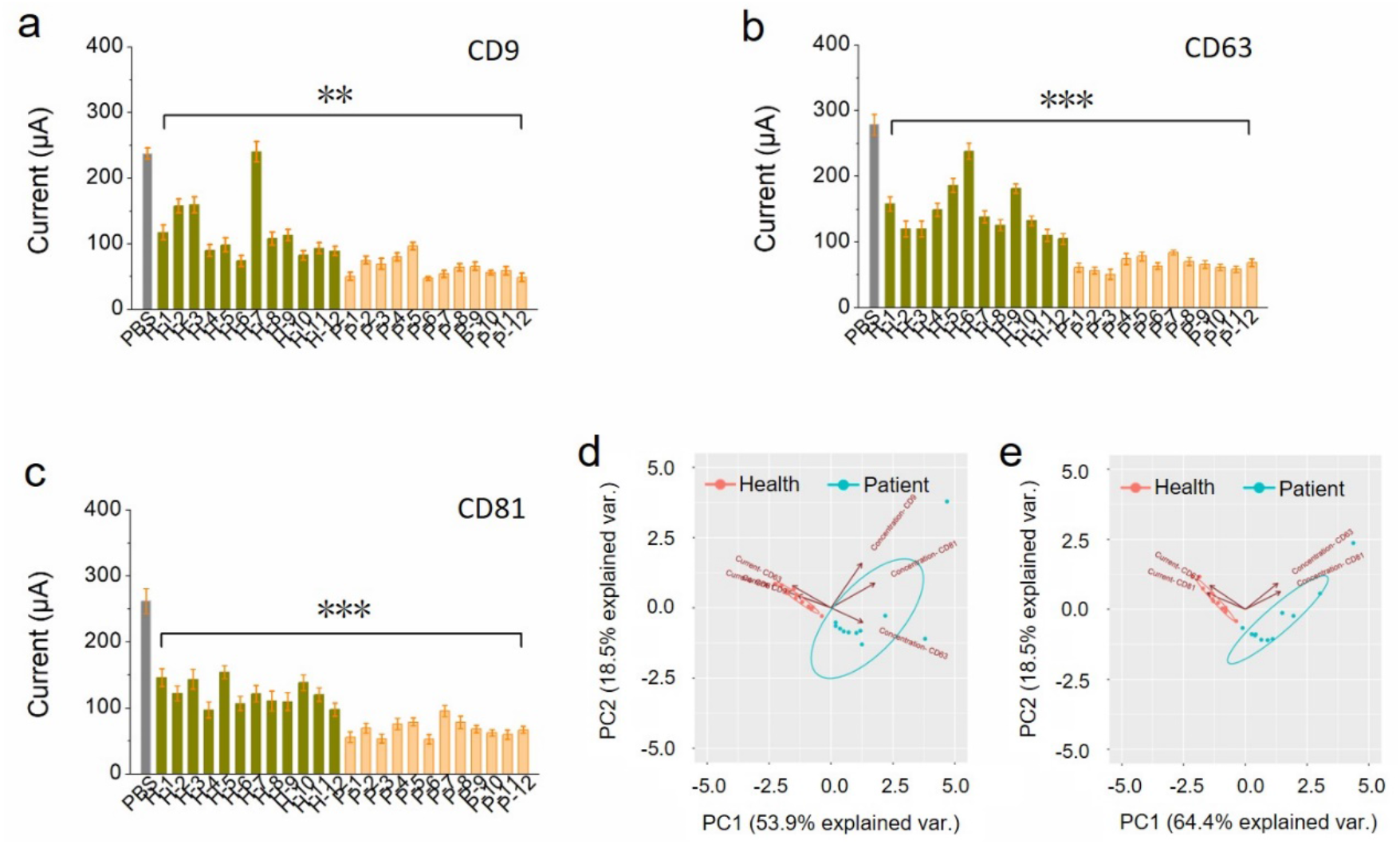
Profiling human lung cancer blood samples through ELB. **a, b, c**, Analysis of exosomes secreted by lung carcinoma cells in clinical human blood samples. Error bars represent mean+/-SD of 3-5 independent experiments. Differences with ^**^P<0.005 and ^***^P<0.0001 were considered statistically significant by T-test. **d, e**, Scores plots of PC1 vs. PC2 produced using data from clinical human blood assay.

To enable accurate data processing, we utilized principal component analysis (PCA), which is a statistical analysis method utilized to investigate major patterns in data sets^34^, to analyze variances in each collected sample and grouping them by similarity. The concentration of exosomes was quantified according to the fitted curves in Figure 4. Figure 5d and 5e present the PCA scores plots of human serum samples. When we selected the protein combination of all three markers, the variance levels in PC1 and PC2 were 53.9% and 18.7%, respectively. However, as the protein combination was defined as CD63 and CD81, the clusters of exosome data could be well-distinguished to cancerous group and health control, demonstrating a robust performance for cancer detection.

## Discussion

Using such rational and bottom-up design approach, we demonstrated a generalizable strategy to encode desirable sensing functions into nano-structured materials. Under the synergistic effect of different components within the hybrid nanostructure, a multiplexed, electrochemical liquid biopsy (ELB) platform was developed and tested with human blood samples, providing accurate clinical insights into cancer severity. Furthermore, the design of this platform technology is modular and logical, therefore serving as a general strategy for the detection of other biomarkers.

Owing to the cost-effectiveness and simplicity of electrochemical sensing comparing with the optical orthologs, electrochemical biosensor represents a promising class of point-of-care systems as exemplified by the gold-standard, commercialized glucose sensor^14,35-44^. Despite significant advances in sensing chemistry^44-52^, challenges remained in electrochemical immunosensors have significantly hindered our capability to assemble an integrated sensing system capable of direct, high-sensitivity and high-accuracy profiling of biomarkers through immunochemistry. First, most of the proteins and nucleic acids are not redox active^42,53^. Therefore, to achieve quantification on an electrochemical surface, characterizing change of surface impedance or constructing redox active molecular interface is required with inevitable introduction of external redox molecules^14,40^. By introducing solid-phase redox active species, we encoded the capability of signal transduction directly onto the electrode materials, providing a generalizable redox-active material interface. Second, surface modification has been significantly investigated and optimized in order to achieve immobilization of recognition elements with optimal conformations^54,55^. By forming homogeneous 3D nanopillars and introducing ZIF-90 with active CHO groups onto the electrode surface, we were able to immobilize conformational ideal and biological active antibody directly onto the electrode without tailoring the surface chemistry, delivering a simple fabrication method for immunosensor. The functionalized nanopillar is also capable of target sorting from complex matrix. Furthermore, the 3D nanostructure built with high-electron conduction materials also provide sufficient electron transfer and anti-fouling properties^11^. Finally, to enable a high-accuracy sensing system or a multi-use sensor, capability of multiplexed analysis is desirable^50,56^. Therefore, multiplexed microfluidic channels were designed to accommodate sensor fabrication and sample input. By examining the overall sensor performance with clinical samples, we confirmed the capability of the rationally engineered ELB platform on profiling and differentiating of physiologically relevant markers. These results indicate that our rational engineering design through encoding sensing function into material interface is a general approach for the developments of electrochemical biosensors.

## Methods

### Synthesis of 2H-MoS_2_

Ultrathin MoS_2_ with 2H-phase was synthesized by sonication exfoliation of bulk MoS_2_ in liquid nitrogen^6^. Typically, 0.1 g MoS_2_ was dispersed into 20 mL KOH (1 g) solution. The mixture was then stirred vigorously for 24 h at 80 °C. After being cooled down to room temperature, liquid nitrogen was quickly poured into the mixture to quench the reaction. The quench process was performed at least four times to effectively prevent the van der Waals forces binding in bulk MoS_2_. The mixture was then sonicated for 3 h and collected by vacuum filtration. The ultrathin 2H-MoS_2_ was finally obtained after drying in vacuum at 60 °C overnight.

### Synthesis of MoS_2_/ZnO/ZIF-90

The synthesis of ZnO@MoS_2_ was carried out through seed-assisted growth strategy. First, the obtained 2H-MoS_2_ was dispersed into ethanol (0.4 mg/mL), and then 1.6 mL of obtained suspension was deposited on a clean silicon wafer with a size of 1 × 1 cm^2^. 250 μL of Zn(Ac)_2_/ethanol solution (5 mM) was then dropped onto 2H-MoS_2_. After drying at room temperature, the silicon wafer was transferred to a tube furnace and calcinated for 20 min at 350 °C. The above silicon wafer loaded with 2H-MoS_2_ and ZnO seeds was laid face down on a solution containing 25 mM Zn(NO)_3_·6H_2_O, 25 mM hexamethylene tetramine, 6 mM polyethylene imine and 80 mL deionized water. The solution was sealed in a 100 mL premium bottle (Schott Duran) and kept at 90 °C for 5 h. After the reaction was completed, the silicon wafer was washed with ethanol and deionized water for three times. ZIF-90 films were grown *in situ* on ZnO@MoS_2_ by using ZnO nanopillars as self-sacrificing template and precursor to directly react with the vapor phase of imidazole-2-formaldehyde. The silicon wafer loaded with MoS_2_/ZnO was suspended in a 100 mL premium bottle with 0.5 g imidazole-2-formaldehyde at the bottom. The vial was kept at 150 °C for 30 min. After the completion of the reaction, the silicon wafer was quickly transferred to a 150 °C vacuum oven and kept for 10 min to remove impurities. The MoS_2_/ZnO/ZIF-90 was finally obtained after cooling down to room temperature.

### Isolation and Concentration Determination of Exosome

Exosomes were harvested from supernatant of human non-small cell lung cancer cell A549 using a QIAGEN Assay kit, Article No. P76064. Briefly, the supernatant was filtered before freezing and then mixed with an equal volume of Buffer XBP evenly and warmed up to room temperature. The supernatant/XBP mixture was added to an exoEasy spin column and centrifuged at 500 g for 1 min. The flow-through fluid was discarded and the column was placed back into the same collection tube. 10 mL buffer XWP was added and centrifuged at 5000 g for 5 min to remove residual buffer. The spin column was then transferred to a fresh collection tube. 400 μL buffer XE was added to the membrane and incubated for 1 min followed by centrifugation at 500 g for 5 min to collect the eluate, the eluate was then re-applied to the exoEasy spin membrane and incubated for 1 min. Centrifugation at 5000 g for 5 min was performed to collect the eluate and the finally extracted exosomes were transferred to an appropriate tube for concentration determination.

The concentration of extracted exosomes was determined using a Beyotime BCA protein Assay Kit, Article No. P0012. An appropriate amount of BCA working solution was prepared according to the instruction and placed at room temperature for 24 h to keep it stable. PBS buffer was used to completely dissolve the protein standard and maintain its concentration at 0.5 mg/mL. A series of increasing volume of standard/PBS solution ranging from 0 μL to 20 μL was added to the standard plate of 96 wells respectively. Additionally, 20 μL of exosomes suspension was added to the sample plate of 96 wells. 200 μL of BCA working solution was added to each well and placed at 37 °C for 30 min. The microplate reader was used to measure the light intensity under 562 nm wavelength light and the concentration of exosomes was calculated according to the standard curve.

### Fabrication of Electrochemical sensor chip

The microfluidic electrochemical sensor chip was configured with a modified five-electrode system and sealed with polydimethylsiloxane (PDMS) layer. The electrochemical electrodes including three working electrodes (WE) as well as their shared reference electrode and counter electrode were screen-printed on polyethylene terephthalate (PET) substrate. Both working electrodes and counter electrode were printed with carbon and silver conductive ink, the reference electrode was printed with silver/silver chloride (Ag/AgCl) ink. The as-prepared MoS_2_/ZnO/ZIF-90 material (4 mg) was dispersed into the mixture of ethanol (100 μL) and Nafion 117 solution (25 μL) forming a homogeneous ink, then locally deposited onto the surface of working electrodes. A 3D-printed template assisted method was used to fabricate PDMS microchannels, and the resin template was printed via a high-resolution DLP/SLA 3D printer (nanoArch S140). The PDMS layer was peeled off from the resin template and then bonded to PET sensor chip by air plasma treatment using a PDC-002-HP Plasma Cleaner (HARRICK PLASMA) under 45 W of treating power in vacuum for 90 s.

### Electrochemical Test

Electrochemical tests were operated on an electrochemical workstation (VSP-300, BioLogic, France). Electrochemical impedance spectroscopy (EIS) was performed in a solution containing 6.7 mM MgCl_2_, 7.6 mM potassium ferrocyanide, 6.7 mM potassium ferricyanide and 300 mL deinoized water with frequency ranging from 10^5^ to 10^−2^ Hz by applying amplitude of 10 mV. The DPV test was performed on the microsensor chip in a potential interval of 0-0.4 V vs. Ag/AgCl and CV test was performed in a potenial interval of -0.6 V-0.0 V and 0.0 V-0.6 V to evaluate the electrochemical response signal.

### Electrochemical immunoassays

Since exosome marker is not exclusive, some of the popular antibodies targeted against exosome associated antigens are cluster of differentiation. Here, three typical exosomes markers (CD9, CD63 and CD81) are employed for the quantification of exosomes in biological samples. In addition, the developed microsensor chip allows multiplex analysis of exosomes from a single sample, which would offer high accuracy and great efficiency through the mutual validation of the acquired data for multiple markers. Firstly, Anti-CD9 antibody/PBS (10 μg/mL), Anti-CD63 antibody/PBS (10 μg/mL), Anti-CD81 antibody/PBS (10 μg/mL) solution were separately injected into different channels through inlet 1, 2 and 3, and then incubated at 4 °C for 12 h. After that, the microchannels were washed three times with wash buffer (PBS+0.05% Tween 20) to remove the unbonded antibodies. In order to perform the assays, the WEs were then blocked for 1h at room temperature with 150 μL of 5% BSA in PBS, and again washed three times with wash buffer. The assays were firstly verified by using recombinant CD (9, 63 and 81) proteins, and then calibrated using culture supernatant from human non-small cell lung cancer cell line A54. All the measurements were repeated at least three times to ensure reproducibility. Clinical blood samples were collected at Tongji Hospital and analyzed in duplicates.

## Acknowledgements

This work was funded by National Key Research and Development Program of China (No. 2021Yfb3200804), Shanghai “Road and Belt” International Cooperation Project (No. 19520744200), Key Basic Research Program of Science and Technology Commission of Shanghai Municipality (20JC1415300). Y.Z. appreciates the financial support of Henan International Joint Research Laboratory of Nanocomposite Sensing Materials. C.L and Y.D appreciate the financial support of Wallace R. Persons Foundation.

## Author contributions

Y.Z. and Y.D. conceptualized the project, interpreted the results, and wrote the manuscript. H.Z. and Z.Y. performed the experiments and analyzed the data. X.G., W.C. and Y.Z. participated in the experiments on isolation and concentration determination of exosomes. All authors revised the manuscript.

## Data and materials availability

All data and materials are available from corresponding author upon reasonable request.

## Supplementary Materials

Materials and Methods, Supplementary Figures

